# E3-specific degrader discovery by dynamic tracing of substrate receptor abundance

**DOI:** 10.1101/2022.10.10.511612

**Authors:** Alexander Hanzl, Eleonora Barone, Sophie Bauer, Hong Yue, Radosław P. Nowak, Elisa Hahn, Eugenia V. Pankevich, Anna Koren, Stefan Kubicek, Eric S. Fischer, Georg E. Winter

**Affiliations:** CeMM Research Center for Molecular Medicine of the Austrian Academy of Sciences, 1090 Vienna, Austria; Department of Cancer Biology, Dana-Farber Cancer Institute, Boston, MA, USA; Department of Biological Chemistry and Molecular Pharmacology, Harvard Medical School, Boston, MA, USA; Proxygen GmbH, 1030 Vienna, Austria

## Abstract

Targeted protein degradation (TPD) is a new pharmacology based on small-molecule degraders that induce proximity between a protein of interest (POI) and an E3 ubiquitin ligase. Of the approximately 600 E3s encoded in the human genome, only around two percent can be co-opted with degraders. This underrepresentation is caused by a paucity of discovery approaches to identify degraders for defined E3s. This hampers a rational expansion of the druggable proteome, and stymies critical advancements in the field, such as tissue- and cell-specific degradation. Here, we focus on dynamic NEDD8 conjugation, a posttranslational, regulatory circuit that controls the activity of 250 cullin RING E3 ligases (CRLs). Leveraging this regulatory layer enabled us to develop a scalable assay to identify compounds that alter the interactome of an E3 of interest by tracing their abundance after pharmacologically induced auto-degradation. Initial validation studies are performed for CRBN and VHL, but proteomics studies indicate broad applicability for many CRLs. Among amenable ligases, we select CRL^DCAF15^ for a proof-of-concept screen, leading to the identification of a novel DCAF15-dependent molecular glue degrader inducing the degradation of RBM23 and RBM39. Together, this strategy empowers the scalable identification of degraders specific to a ligase of interest.

## Introduction

Modulation of proximity between macromolecules is a central regulatory layer in most cellular processes. Chemically inducing proximity between two target proteins is an established mechanism of action of natural products as well as synthetic small molecules, and has expanded to clinical applications^1–3^. In some instances, drug-induced proximity results in a functional inhibition of one participating protein via steric confinements imposed by the other effector. In other instances, the result of induced proximity can be an emerging effect where one protein partner is endowed with a novel, *neomorphic* function^4^. This scenario is best illustrated by small-molecule degraders that recruit proteins of interest (POIs) to an E3 ligase. The change in E3 ligase interactome consequently leads to ubiquitination and degradation of the POI, which it would not recognize in absence of the small molecule^5^. This concept of targeted protein degradation (TPD) comes with several advantages compared to inhibitor-centric approaches^6^. However, one of its current major limitations is the reliance on only a small set of amenable E3 ligases^7^. Existing degrader modalities suggest that members of the cullin RING ligase (CRL) family are particularly compatible with TPD strategies. Approximately 250 distinct CRL ligases are encoded in the human genome, organized around one of seven cullin scaffolding proteins^8,9^. CRLs are highly dynamically regulated and often assembled and actively decommissioned based on substrate availability^10,11^. Both processes are dependent on the deposition and removal of the small ubiquitin-like modifier NEDD8^12^. Attachment of NEDD8 on the cullin backbone stabilizes an active CRL complex primed for ubiquitination of a substrate, enhancing its catalytic ability up to 2000-fold^13^. Conversely, “de-neddylation” from the cullin backbone by the COP9 signalosome (CSN)^14^ allows the exchange of substrate recruiting receptors (SRs) of CRLs, thereby reshaping the ubiquitinated proteome in a cell^15^. In absence of continuous substrate supply, this enables adaptation to cellular stimuli. If CRL decommissioning is perturbed, for instance by a dysfunctional CSN, the respective CRL is hence arrested in a continuous state of activity. In the absence of substrate, this eventually leads to a process termed “auto-degradation” where the CRL-associated E2 ubiquitinates the substrate receptor^16,17^.

Only a few CRLs have thus far been harnessed for TPD, primarily via so-called hetero-bifunctional proteolysis targeting chimeras (PROTACs)^18^. These degraders bind the E3 ligase and the POI with distinct chemical moieties connected by a linker. The modular design of PROTACs allows facile chemical and thereby neo-substrate alteration. However, their degradable proteomic space is limited to ligandable targets. Molecular glue degraders (MGDs), on the other hand, are monovalent small molecules that stabilize a recognition surface between the ligase and the POI via degrader-protein and protein-protein interactions (PPIs). Mechanistic dissection of the clinically approved immunomodulatory drugs (IMiDs) has unveiled such a mechanism for the CRL4^CRBN^ dependent degradation of zinc finger transcription factors^19–21^. Similar studies have further identified MG degraders of splicing^22,23^ and translation factors^24^ via a limited set of E3 ligases also including CRL^DCAF15^.

Recent advances in chemo-proteomics workflows have augmented the identification of E3 ligase binders conducive to PROTAC development^25–29^. However, this has not yet led to the identification of novel MGDs. Rational MGD discovery would benefit from a technology that measures drug-induced changes in the E3 ligase interactome in an unbiased fashion. Methods yielding such proteome-wide interaction data however lack the required throughput. At the same time, prospective high throughput degrader discovery has traditionally been confined to readouts covering pre-defined ligase-substrate pairs using recombinant proteins^30^.

Here, we leverage the unique regulatory dynamics of CRLs to design a scalable assay informing on drug-induced changes to the interactome of a predefined ligase of interest. We find that degrader-mediated neo-substrate recruitment to a CRL rescues its SR from experimentally induced auto-degradation. Dynamic SR “tracing” via life-cell bioluminescence quantification allows degrader screening in a target agnostic, scalable and time-resolved manner. We first benchmark this assay with known PROTACs and MG degraders targeting CRL4^CRBN^ and CRL2^VHL^. Next, we determine the number of E3 ligases amendable to this approach by measuring substrate receptor auto-degradation upon CSN inhibition. Among the destabilized ligases, we select CRL^DCAF15^ for performing in-depth mechanistic studies and a proof-of-concept screen with a bespoke library of 10’000 compounds. During hit validation, the compound dRRM-1 proved to be a DCAF15-dependent, chemically distinct molecular glue degrader of RBM39 and RBM23. Taken together, the presented technology empowers the scalable identification of molecular glue degraders specific to a ligase of interest.

## Results

### E3 ligase abundance serves as a proxy for neo-substrate recruitment to E3 ligases

CRL activity has been implicated in degrader potency and used for the identification of novel MGDs.^31,32^ The current approaches however are limited to targets essential for cellular viability and don’t allow ligase focused discovery efforts. Active CRLs, in absence of their substrate can ubiquitinate their own substrate receptor in a process termed auto-degradation^17,33,34^. This fundamental mechanism has been implicated in CRL adaptation to substrate availability and cellular stimuli^10,11^. Here, we envisioned that chemically re-establishing substrate availability by degrader-mediated recruitment of a neo-substrate to a constitutively active, auto-degrading CRL should stabilize the ligase. Ubiquitination will be deflected from the SR to the neo-substate and consequently increase the SR abundance (**Figure 1A**). To test this hypothesis, we treated the near haploid chronic myeloid leukemia cell line HAP1 with the CRL4^CRBN^ dependent GSPT1 molecular glue degrader CC-885.^24^ Indeed, we observed that CC-885 treatment rescued CRBN autodegradation (**Figure 1B**). However, based on the minor increase compared the vehicle (DMSO) treatment, we surmised that steady-state CRBN auto-degradation has only a minor contribution to its turnover (compare CRBN levels in lanes 1 and 2). Given that cullin scaffold engagement of each of the ~250 SRs varies greatly^10,11^, also their auto-degradation will depend on factors such as cell type and state. We therefore reasoned that enrichment of the pool of constitutively active CRLs allows the augmentation the dynamic range of this observation (**Figure 1A**). NEDD8 is the central post-translational modification governing CRL activity^13^ and treatment with the de-neddylation inhibitor CSN5i-3 was previously shown to lock CRLs in a state of constitutive activation that leads to auto-degradation^34,35^. Indeed, treatment of HAP1 cells with 500 nM CSN5i-3 yielded a significant destabilization of the CRBN substrate receptor. As hypothesized, recruitment of GSPT1 via CC-885 rescued the induced auto-degradation almost to DMSO treated levels (**Figure 1B**)^24^. This striking response to molecular glue treatment led us to next explore possibilities to develop a discovery approach where novel degraders are identified by their ability to rescue a CRL from a constant state of autodegradation. Compared to existing methods, this strategy would enable the identification of degraders specifically for an *a priori* defined ligase. Moreover, it is not restricted to POIs that are essential for cellular fitness or are otherwise implied in any measurable cellular phenotype. To make this concept amenable to degrader-discovery at scale, we next sought to advance this approach towards scalable protein abundance measurement across many timepoints upon drug treatment.

**Figure 1.**
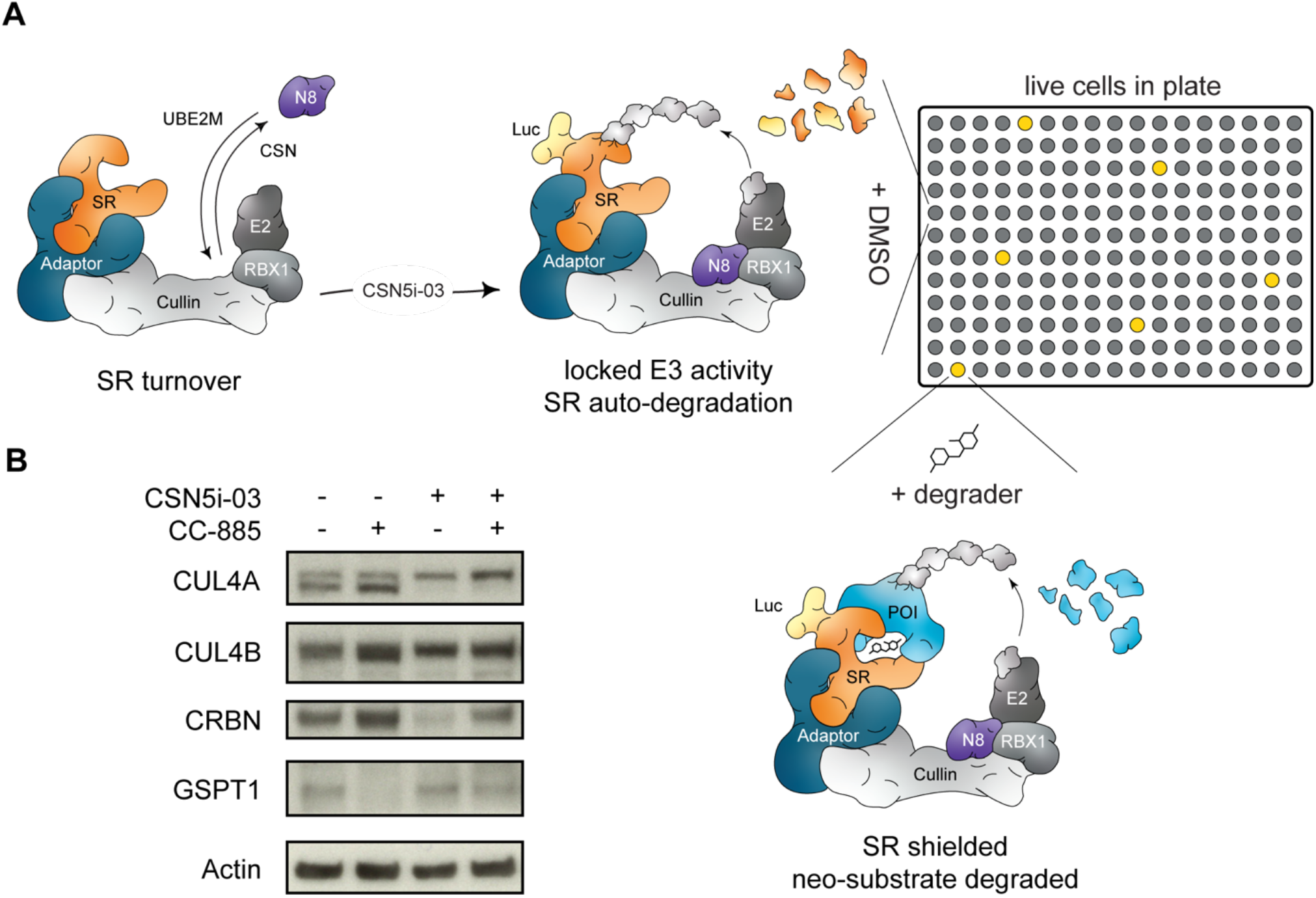
Degrader-induced neosubstrate recruitment rescues CRBN from auto-degradation. (A) Schematic depiction of the ligase tracing approach. Cullin RING ligase decommissioning is mediated through deposition of NEDD8 (N8) on the cullin backbone via the COP9 Signalosome (CSN). Inhibition of the CSN locks cullin ligases in an active conformation leading to auto-ubiquitination and -degradation of the luciferase tagged substrate receptor. In 96- and 384-well plates, addition of a degrader compound can shield the substrate receptor and rescue its auto-degradation leading to detection of luciferase signal. (B) Protein levels in KBM7 WT cells pre-treated for 10 min with DMSO or CC-885 (100 nM) followed by treatment with DMSO or CSN5i-3 (500 nM) for 4 h. Representative images of *n* = 2 experiments.

### Many CRLs are amenable to dynamic tracing of E3 ligase abundance

We next proceeded to validate and benchmark the idea of “ligase tracing” for degrader identification with CRL2^VHL^, a ligase that has often been employed for targeted protein degradation via PROTACs^36,37^. Generating VHL-NanoLuc knock-in HAP1 cells, allowed us to measure VHL abundance in lytic measurements in 384-well plate format. Upon induction of auto-degradation via CSN5i-3 treatment, VHL was destabilized within hours (**Figure 2A**). In live-cell measurements we further found this to be a dose- and time-dependent process, strengthening the causal link between CRL activity and SR auto-degradation (**Figure S1A**). Consistent with our hypothesis, co-treatment with the BET PROTACs ARV-771 and MZ1 profoundly rescued auto-degradation of endogenous VHL (**Figure 2A** and **Figure S1A**)^38,39^. Overexpression of NanoLuc-tagged VHL further validated these results where BET PROTACs showed dramatic VHL stabilization (data is normalized to the auto-degraded state induced by CSN5i-3 treatment, **Figure 2B**). Live-cell measurements also allowed us to compare responses of active BET PROTACs to inactive negative controls, and to further expand the target space to additional substrates. In line with the proposed mechanism, the enantiomer *cis*-MZ1 which is deficient for VHL binding did not elicit a response in our ligase tracing (**Figure 2B**). This was also validated in assays performed with NanoLuc-VHL knock-in cells (**Figure S1A**). Similarly, the SMARCA2/4 degrading PROTAC^40^ ACBI1 showed a pattern where only the active compound provoked changes in the auto-degradation behavior of VHL, while the enantiomer that is deficient for VHL binding failed to stabilize VHL (**Figure 2B**). Next, we excluded that VHL stabilization is prompted simply by small-molecule binding to VHL. Indeed, no stabilization was observed with the VHL ligand VH-032 (**Figure 2B** and **Figure S1A**). In sum, these chemical controls validate the dependency of ligase tracing on ternary complex formation and dual target engagement for a positive stabilization signal. Overall, this indicates that CRL substrate receptor abundance can be used as a proxy for degrader-induced neosubstrate recruitment.

**Figure 2.**
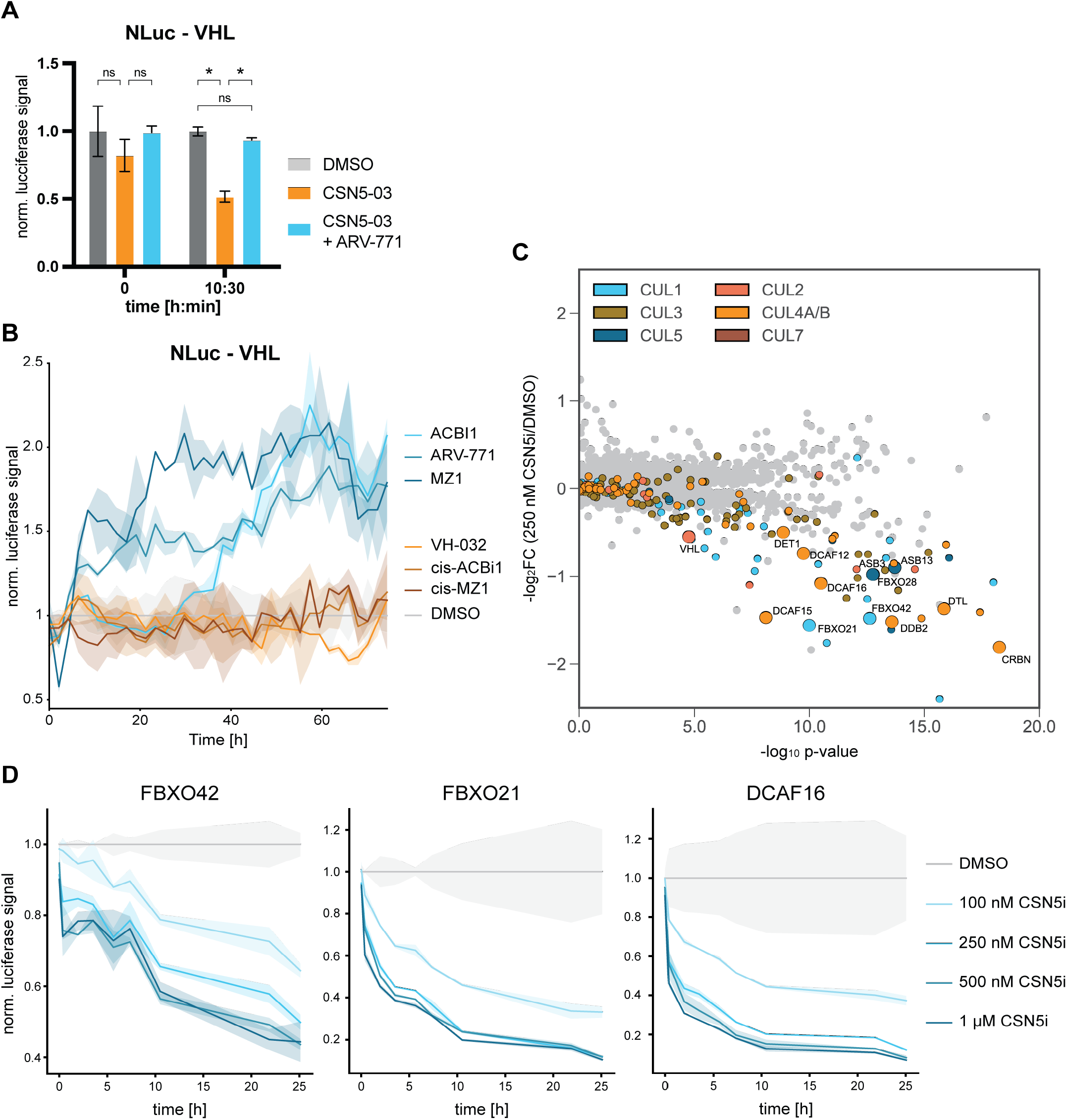
Mapping E3 ligases amendable to ligase tracing. (A) Lytic luciferase measurement of HAP1 VHL-NanoLuc knock-in cells at the indicated timepoints after treatment with DMSO, CSN5i-3 (100 nM) or CSN5i-3/ARV771 co-treatment (100 nM & 500 nM respectively). Luciferase signal is normalized to DMSO treatment at each timepoint. Mean ±s.d. of *n* = 2 replicates. (B) Live-cell luciferase measurement of HAP1 VHL-NanoLuc expressing cells treated with CSN5i-3 (100 nM) or CSN5i-3 and the indicated degrader (100 nM & 500 nM respectively). Luciferase signal is normalized to DMSO treatment at each timepoint. Mean ± s.d. of *n* = 2 replicates. Representative data of *n* = 2 experiments. (C) Volcano plot depicting global log_2_-fold changes of protein abundance in HAP1 cells treated with CSN5i-3 (250 nM) for 8 h. CRL substrate receptors are labeled in the indicated colors. SRs selected for validation via luciferase tagging are highlighted. Data of *n* = 3 replicates. (D) DMSO normalized live-cell luciferase signal of HAP1 cells harboring endogenous NanoLuc knock-ins for the indicated SRs. Cells were treated with DMSO or CSN5i-3 at indicated concentrations and measured over time. Mean ± s.d. of *n* = 2 replicates. Statistical analysis via a two-tailed *t*-test (α = 0.05), \**P* < 0.05.

CRL2^VHL^ and CRL4^CRBN^ are very well characterized E3 ligases in TPD. To significantly expand the space of degraders and neo-substrates we next set out to assay which CRLs are amendable to our ligase tracing assay. As the active CRL pool is shaped to the particular cellular needs at any given time, this is likely cell-line and -state specific. We chose to perform global proteomics in HAP1 cells, given that its near-haploid karyotype facilitates endogenous tagging of SRs with NanoLuc. Expression proteomics experiments were recorded after 250 nM and 1 μM CSN5i-3 treatment for 8 hours (**Figure 2C** and **Figure S1B**). In line with our hypothesis, most of the destabilized proteins were cullin associated substrate receptors. Among these destabilized CRLs we selected twelve SRs for either a NanoLuc knock-in or overexpression strategy in HAP1 cells. Measuring the SR abundance upon CSN5i-3 induced auto-degradation resulted in dose- and time dependent destabilization (**Figure 2D** and **Figure S1C**). This mimics our previous results and highlights that the ligase tracing approach can be expanded to many CRLs.

### Ligase tracing screen identifies a novel DCAF15-dependent molecular glue degrader

Discovery of novel molecular glue degraders has historically been driven by chance. After benchmarking the ligase tracing strategy with known degraders, we next set out to validate it in a chemical screening approach. To this goal, we chose to adopt the ligase tracing approach for CRL4^DCAF15^. DCAF15 can be targeted by aryl sulfonamides such as indisulam to recruit and ubiquitinate the splicing factor RBM39, hence it enabled us to assemble a bespoke chemical library around a chemotype know to engage DCAF15^22,23^. Additionally, our previous results suggested that DCAF15 is strongly auto-degraded upon CSN inhibition, thus making it eligible to ligase tracing (**Figure 2C** and **Figure S1C**).

To measure DCAF15 abundance we initially tagged endogenous *DCAF15* with the split luciferase eleven amino acid peptide HiBit in HEK-293 cells supplemented with the complimentary NanoLuc part LgBit.^41,42^ Upon treatment with indisulam, we observed an expected rescue of CSN5i-3-induced DCAF15 auto-degradation (**Figure S2A**). Due to low expression levels of endogenous *DCAF15,* we however decided to proceed with overexpression of HiBit-DCAF15 in *DCAF15^-/-^* cells. In this system we encountered profound SR stabilization upon indisulam treatment even in the absence of CSN inhibition, likely due to a strong cellular auto-degradation in response to the SR overexpression (**Figure 3A**). Next, we proceeded to determine DCAF15 abundance via lytic endpoint measurements. Validating our Western blot results, we observed destabilization upon CSN5i-3 treatment and profound stabilization of HiBit-DCAF15 by indisulam treatment (**Figure 3B**). Furthermore, we could reproduce this stabilization also in live-cell measurements, and could also extend it to another, previously identified RBM39 molecular glue degrader (dCeMM1) (**Figure 3C** and **Figure S2B)** ^32^. Next, we set out to test whether indisulam mediated stabilization was specific to DCAF15 by performing ligase tracing in a variety tagged NanoLuc -SR cells. Indeed, rescue of ligase degradation was specific to HiBit-DCAF15 cells, while DCAF16, FBXO21, VHL and FBXO42 NanoLuc knock-ins remained unchanged (**Figure S2C**). Importantly, the increase in DCAF15 abundance was not driven through changes in RNA expression as exemplified by *DCAF15* qPCR (**Figure S2D)**.

**Figure 3.**
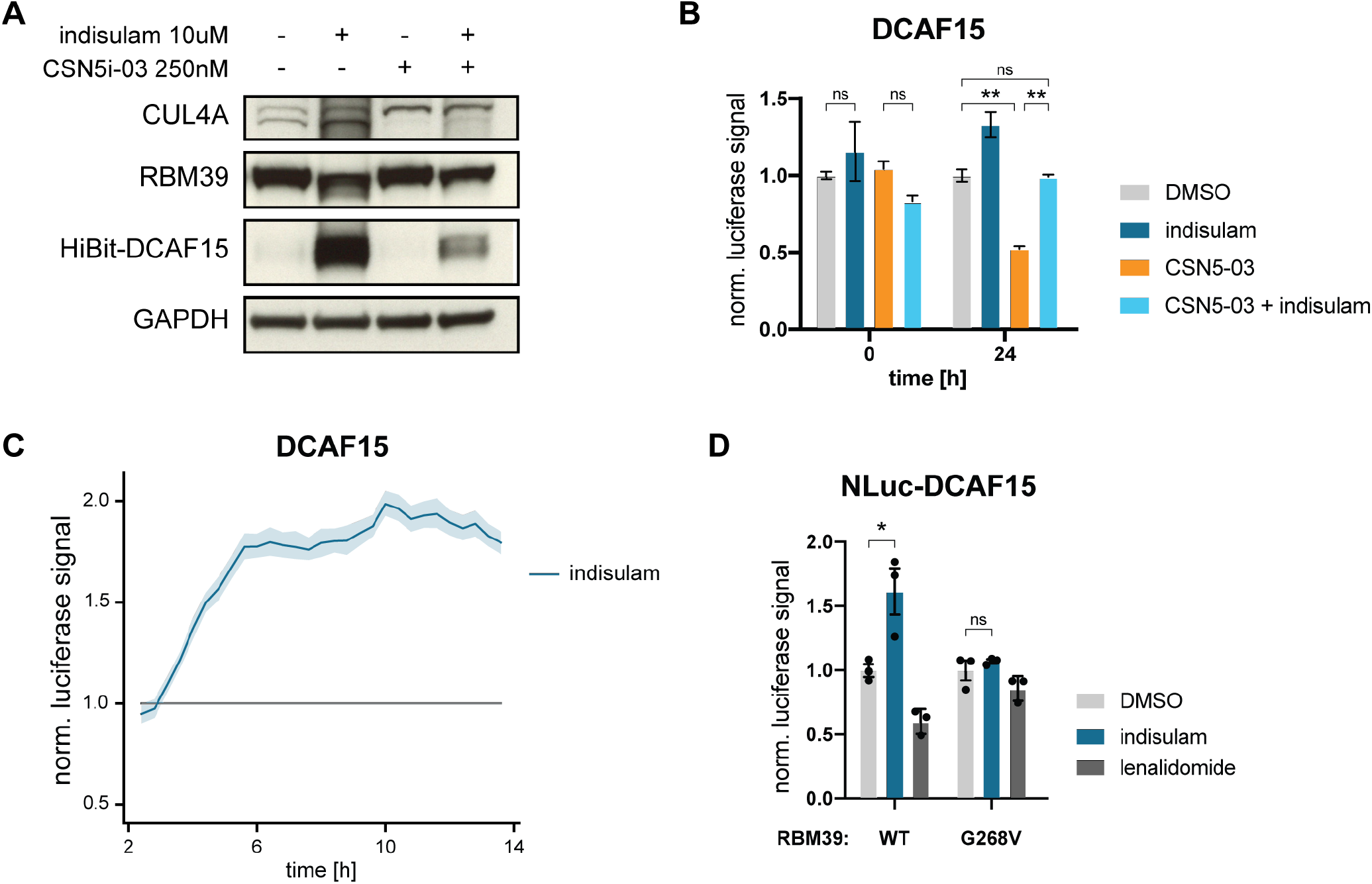
Ligase tracing detects molecular glue degrader dependent DCAF15 stabilization. (A) Protein levels in HEK293t DCAF15^-/-^ cells with reconstitution of HiBit-DCAF15 treated with indisulam (10 μM) or CSN5i-3 (250 nM) for 24 hrs. Representative images of *n* = 2 experiments. (B) Bar graph depicting DMSO normalized lytic luciferase signal of HEK293t DCAF15^-/-^ cells with reconstitution of HiBit-DCAF15 + LgBit measured at the indicated timepoints after treatment with DMSO, indisulam (10 μM), CSN5i-3 (250 nM) or CSN5i-3/indisulam co-treatment (250 nM & 10 μM respectively). Mean ±s.d. of *n* = 2 replicates. Representative data of *n* = 2 experiments. (C) DMSO normalized live-cell luciferase signal of HEK293t DCAF15^-/-^ cells with reconstitution of HiBit-DCAF15 + LgBit treated with indisulam (10 μM) or DMSO. Mean ±s.d. of *n* = 3 replicates. Representative data of *n* = 3 experiments. (D) Bar graph depicting DMSO normalized live cell luciferase signal of HAP1 WT and RBM39^G268V^ cells with ectopic expression of HiBit-DCAF15 + LgBit measured 32 h after treatment with DMSO, indisulam (10 μM) or lenalidomide (10 μM). Mean ±s.d. of n = 3 replicates. Representative data of *n* = 2 experiments. Statistical analysis via a two-tailed *t*-test (α = 0.05), \**P* < 0.05, \*\**P* < 0.01.

In its initial identification, RBM39 degradation by indisulam was shown to be dependent on the glycine residue 268 in RBM39^22^. Mutation of this amino acid to a valine abrogated neo-substrate recruitment and degradation^43–45^. This critical dependency on a given protein surface topology enabled us to genetically validate the observed DCAF15 stabilization is indeed caused by functional neo-substrate recruitment. To this end, we endogenously engineered a RBM39^G268V^ mutation in near-haploid HAP1 cells overexpressing HiBit-DCAF15 (**Figure 3D**). Indeed, indisulam only induced a stabilization effect in the RBM39^WT^ cells while no change could be detected in a RBM39^G268V^ background. Of note, the CRL4^CRBN^ molecular glue degrader lenalidomide did not show any stabilization effect (**Figure 3D**). Together this indicates that stabilization of DCAF15 with different RBM39 degraders is dependent on drug-induced neo-substrate recruitment. Furthermore, this highlights how CRL4^DCAF15^ presents a viable system for molecular glue degrader identification via our ligase tracing approach.

Having established live-cell ligase tracing for CRL4^DCAF15^, we next set out to determine this assay’s viability for degrader discovery screening. To this end we assembled a library of 10,000 sulfonamides of which ~ 8,000 were arylsulfonamides leveraging the known MG chemical space for DCAF15. Ligase tracing for DCAF15 with this library showed a profound stabilization of SR levels with positive controls (indisulam, dCeMM1) in concordance with previous results (**Figure 4A**). As this assay harbors the inherent advantages of gain-of-signal approaches^46^ we found a surprisingly low hit rate among other sulfonamides. In fact, similar stabilization effects were only observed for the aryl sulfonamide we termed dRRM-1, which shared some structural similarity to indisulam and dCeMM1 (**Figure 4A** and **Figure 4B**). Given this similarity, we determined RBM39 levels upon cellular treatment with dRRM-1 and detected DCAF15 dependent degradation similar to indisulam and dCeMM1 (**Figure 4C**). Upon docking of dRRM-1 to a published crystal structure of DDB1ΔB-DDA1-DCAF15-E7820-RBM39, we further identified a shared binding mode with the previously described sulfonamide degrader E7820, suggesting a similar mode-of-action via CRL4^DCAF15^ mediated degradation of RBM39 (**Figure 4D**)^43–45^. We could further validate induced degradation via C-terminal knock-in of HiBit to *RBM39* and measuring its abundance via live-cell luciferase detection (**Figure S3A**). Similarly to known RBM39 degraders such as tasisulam, measurement of E7820 displacement from DCAF15 by dRRM-1 in a TR-FRET assay, revealed a comparatively low binding affinity to DCAF15 (**Figure 4E**)^45^. This however highlights how ligase tracing can detect functionally relevant glue degraders, even though they have comparatively low ligase affinity. Global proteomics experiments revealed that not only RBM39 is degraded via dRRM-1 treatment but also the closely related splicing factor RBM23 (**Figure 4F**). RBM23 shares a high sequence similarity to RBM39 and has previously been shown to be targeted by other sulfonamides^45,47^. Intrigued by the potential degradation of RBM23 over RBM39 by dRRM-1, we generated a C-terminal RBM23-NanoLuc knock-in HAP1 cell line and measured its abundance upon sulfonamide treatment. Indisulam and dRRM-1 led to similar time dependent RBM23 degradation, which could be rescued by co-treatment with the proteasome inhibitor carfilzomib (**Figure 4G**).

**Figure 4.**
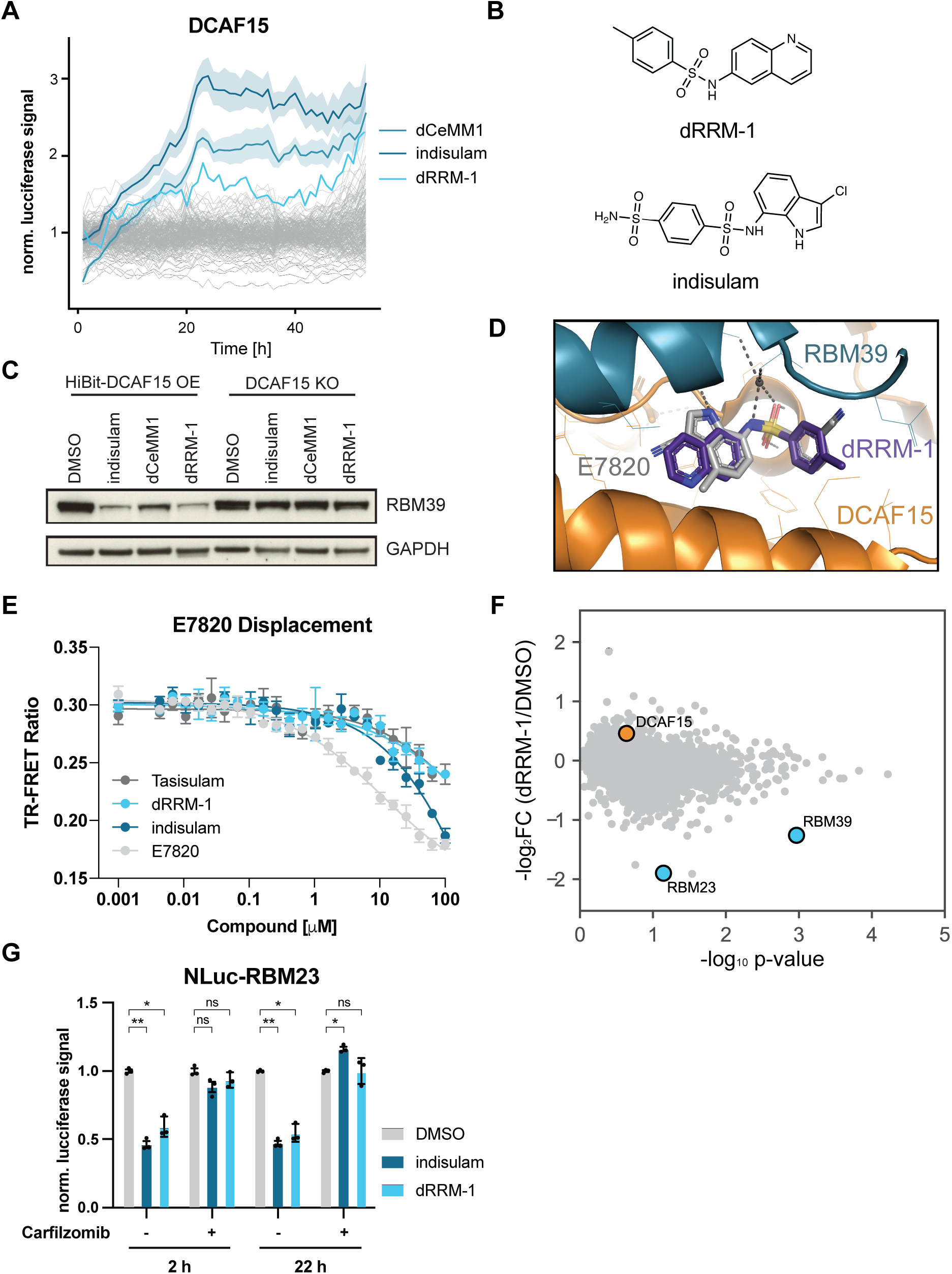
A ligase tracing screen for DCAF15 MGDs identifies dRRM-1. (A) DMSO normalized live-cell luciferase signal of HEK293t DCAF15^-/-^ cells with reconstitution of HiBit-DCAF15 + LgBit treated with 10 μM control compounds (indisulam, dRRM-1 or DMSO) or screening compounds (10 μM each, 200 compounds shown). For indisulam and dCeMM1 data represents mean ±s.d. of all measured wells. (B) Chemical structures of dRRM-1 and indisulam. (C) Overlay of molecular docking of dRRM-1 in the crystal structure of DDB1ΔB-DDA1-DCAF15:E7820:RBM39 (PDB: 6Q0R). (D) Protein levels in HEK293t DCAF15^-/-^ cells with reconstitution of HiBit-DCAF15 treated with indisulam, dCeMM1 or dRRM-1 for 10 hrs. Representative images of *n* = 2 independent experiments. (E) TR-FRET ratio of BODIPY FL-E7820 displacement from biotinylated, terbium labeled, Strep-II-Avi-tagged DCAF15 with increasing amounts of tasisulam, dRRM-1, indisulam or positive control E7820. The emission ratio of 520 nm (BODIPY FL) over 490 nm (terbium) is calculated and depicted as mean ± s.d. from *n* = 3 replicates. (F) Volcano plot depicting global log_2_-fold changes of protein abundance in HEK293t DCAF15^-/-^ cells with ectopic expression of HiBit-DCAF15 and LgBit treated with dRRM-1 (10 μM) for 10 hrs. Data of *n* = 2 independent measurements. (G) Bar graph depicting DMSO normalized live cell luciferase signal of HAP1 RBM23-NanoLuc knock in cells measured at the indicated timepoints after treatment with DMSO, indisulam (10 μM) or dRRM-1 (10 μM). Mean ± s.d. of *n* = 3 replicates. Statistical analysis via a two-tailed *t*-test (α = 0.05), \**P* < 0.001, \*\**P* < 0.0001.

In summary, we outline and validate a CRL-centric phenotypic screening approach that allowed us to identify dRRM-1, a previously undescribed, DCAF15-dependent MGD which retains a binding mode similar to previously described, and serendipitously identified degraders.

## Discussion

TPD promises a paradigm shift in drug discovery by overcoming limitations of inhibitor-centric, occupancy-driven pharmacology through its catalytic components. In addition to expanding the druggable proteome, areas of interest include tissue- and cell-state selective degradation of disease relevant proteins. Delivering on these promises is currently severely limited as only two percent of E3 ligases are amenable to TPD. Methods based on chemo-proteomics have led to the discovery of covalent E3 ligase binders which in turn have spurred PROTAC development^25–29^

Here we outlined a strategy of measuring drug induced changes to the interactome of an E3 ligase of choice by leveraging the regulatory circuits of cullin RING ligases. We benchmark this scalable assay with the two best-studied E3 ligases in TPD, CRL4^CRBN^ and CRL2^VHL^. By use of the CRL2^VHL^ binding ligand VH-032 we show that drug-ligase engagement is insufficient to rescue a ligase from auto-degradation. Instead, the ligase tracing assay specifically reports on drug-induced neo-substrate recruitment. We further profile all E3 ligases amendable to this approach in our given cell line model and choose to perform in-depth studies with CRL4^DCAF15^ given its pronounced auto-degradation and the availability of aryl sulfonamides as a known chemotype potentially capable of co-opting DCAF15. A single point mutation abrogating MGD dependent recruitment of RBM39 to CRL^DCAF15^ was sufficient to disrupt ligase tracing signal highlighting the assay specificity. Among 10,000 sulfonamides tested, we identified dRRM-1 a molecular glue degrader of RBM39 and RBM23 and validate its mode of action via TR-FRET and global proteomics. We conclude that our ligase tracing assay allows identification of functional degrader molecules in an E3 ligase selective but target agnostic way.

This allows selection of therapeutically enticing CRL E3 ligases, taking into account their characteristics such as disease relevance, expression pattern and target complementarity. In fact, recently we have shown that essentiality of an E3 ligase can have profound impact on emergence of resistance to degrader modalities further highlighting the need to expand the targetable E3 ligase space^48^. In principle, ligase tracing assays capture POI recruitment on a proteome-wide scale. Future research will be required to determine thresholds of neosubstrates abundance required for this assay. Of note, the explored neo-substate space can likely be biased by overexpressing pools of targets of interest. An advantage of ligase tracing over other previously reported methods for molecular glue discovery lies in its independence from the neo-substrate’s essentiality status. While discovery of cyclin K molecular glue degraders hinged on their cytotoxicity, the here presented method directly reports on changes to the interactome of an E3 ligase. The general gain-of-signal design of ligase tracing also harbors advantages such as increasing rates of true-positives. Moreover, it is conceivable that drug-indued recruitment of different neo-substrates might prompt different kinetics of ligase restabilization, based on the steric considerations and abundance of a given POI. While future research will be needed to unequivocally validate this hypothesis, ligase tracing of VHL has indicated differentiated stabilization curves for BET PROTACs compared to the SMARCA2/4 PROTAC ACBI1. Finally, one can envision a multiplexing strategy by measuring the abundance of several E3 ligases in the same well via fluorescent protein tagging coupled to a microscopy

Of note, a gain of signal in ligase tracing not necessarily requires neo-substrate degradation but could also be caused by disassembly of the specific ligase, by inhibition of the associated E2, or by even more pleiotropic UPS perturbations. However, such false positives would be identified for most assayed ligases and can hence easily be eliminated. Overall, we believe that the outlined method can be easily adopted to other E3 ligases of interest and facilitate the *de novo* identification of E3 ligase binders and molecular glue degraders in a target agnostic fashion.

## Author contribution

A.H. and G.E.W. conceptualized this study. A.H. with the help of A.K. and S.K. designed and performed E3 ligase luciferase measurements. A.H., S.B., E.B., E.H. and E.V.P. generated over-expression and knock-in cell lines and conducted immunoblotting and cellular drug sensitivity assays. A.H. generated samples for proteomics experiments and visualized resulting data. H.Y. performed TR-FRET assays. R.P.N. performed molecular docking studies. E.S.F., S.K. and G.E.W. supervised the work. A.H. generated figures with input from all authors. A.H. and G.E.W. wrote the manuscript with input from all authors.

## Acknowledgements

We thank the proteomics facility at CeMM (in particular Frédéric Fontaine and André Müller) for assistance with proteomics experiments. CeMM and the Winter laboratory are supported by the Austrian Academy of Sciences. The Winter lab is further supported by funding from the European Research Council (ERC) under the European Union’s Horizon 2020 research and innovation program (grant agreement 851478), as well as by funding from the Austrian Science Fund (FWF, projects P32125, P31690 and P7909).

## Materials and Methods

### Cell lines, tissue culture and lentivirus production

KBM7 cells (a gift from T. Brummelkamp) were grown in IMDM supplemented with 10% Fetal Bovine Serum and 1% penicillin/streptomycin (100 units/ml Penicillin, 100 μg/ml Streptomycin; pen/strep). HAP1 cells were cultured in IMDM supplemented with 10% Fetal Bovine Serum and 1% pen/strep. HEK293T cells (a gift by the Bradner Lab) were grown in in DMEM supplemented with 10% Fetal Bovine Serum and 1% pen/strep. HCT116 cells (a gift by the Superti-Furga Lab) were cultured in DMEM supplemented with 10% Fetal Bovine Serum and 1% pen/strep. pSpCas9(BB)-2A-GFP (PX458) or pSpCas9(BB)-2A-Puro (PX459) was obtained through Addgene (48138 and 62988) and used to transiently express sgRNA against CRBN, VHL, DCAF15 and other genes for knock-out and knock-in generation (sgRNA sequences are summarized in Supplementary Table 1). Clones were single cell seeded and checked for gene deletion via PCR on gDNA or Western blotting. For lentiviral production, 293T cells were seeded in 10 cm dishes and transfected at approx. 80 % confluency with 4 μg target vector, 2 μg pMD2.G (Addgene 12259) and 1 μg psPAX2 (Addgene 12260) using PEI and following standard protocol. The viral supernatant was harvested 72 h after transfection and filtered with a 0.45-μm syringe filter to remove cell debris. Lentivirus was then aliquoted and stored at – 80 °C until transduction of 1 x 10^6^ cells in 1 ml of media plus virus in 24 well plates with the addition of 8 μg per ml polybrene (Sigma) and spin inoculation for 1 h at 2,000 r.p.m. Antibiotic selection was performed 24 to 48 h after transduction with 10 μg ml^-1^ blasticidin.

### Plasmids and cloning

All plasmids used in this study are summarized in Supplementary Table 1. For ectopic expression of E3 ligases tagged with full length NanoLuc, cDNA of the specific genes was ordered in pENTR223 vectors from the BCCM/Belspo consortium as part of the human ORFome library^49^. E3 ligase cDNAs were then cloned into pLenti6.2-ccdB-Nanoluc (Addgene 87075) via gateway LR-recombination cloning (Invitrogen) following manufacturers recommendations.

For cloning of sgRNA cutting plasmids to generate endogenous NanoLuc and HiBit knock-ins, we utilized a universal pX330A_sgX_sgPITCh cutting plasmid via adaptation of a published protocol^50^. sgRNAs targeting the endogenous locus (Supplementary Table 1) were selected to lie as close as possible to the start- or stop codon with the minimal predicted off-target activity. They were introduced to the vector via oligonucleotide annealing and subsequent BbsI-mediated restriction cloning. The second part of this micro-homology mediated knock-in strategy was introduced by adapting the pCRIS-PITChv2 repair template plasmid to contain a N-terminal blasticidin-P2A-2xHA-NanoLuc cassette which was generated via a geneblock and PCR of the flanking PITCh sgRNA target sites. This PCR product was then introduced in MluI linearized pCRIS-PITChv2 vector via NEBuilder 2× HiFi assembly (New England Biolabs). Primers containing 20 to 22 bp homology regions corresponding to the genomic locus 5’ and 3’ of the sgRNA cleavage were used to PCR this cassette. The resulting repair template introducing a blasticidin marker, and a double HA tagged NanoLuc to the genomic locus was reintroduced into MluI linearized pCRIS-PITChv2 vectro backbone with NEBuilder 2× HiFi assembly (New England Biolabs)^51^.

### Endogenous genome editing for knock-in and mutant generation

To generate cell lines expressing HiBit or NanoLuc tagged POIs, HAP1 WT or HCT116 WT cells were seeded into 6-well plates to obtain approximately 70 % confluency the next day. For microhomology mediated knock-in, in each well, PITCh sgRNA/Cas9 and repair template plasmids were transfected at 1.5 μg each via PEI following standard protocol (see Supplementary Table 1 for sgRNA and microhomology sequences). The next day, each condition was split to a 10 cm dish and antibiotic selection for successful editing was started 48 hours after transfection. Single cell selection was ensured by limited dilution into 384 well-plates (seeding at 0.2–1 cells per well in 50 μl) or by picking single colonies directly of the plate. Successful knock-in was characterized via immunoblotting for the introduced HA-tag and/or via genotyping by PCR of the targeted genomic region.

For knock-in of the shorter HiBit-tag in the C-terminus of RBM39 in HCT116, a similar approach was used only that the repair cassette could be introduced via annealed oligos instead of an entire plasmid. For this oligos harboring the 33 bp HiBit-tag flanked by 20 bp homologies from the genomic sgRNA cut site were ordered and co-transfected as described above. Cells were pulse-selected from 24 h post transfection to 72 hrs post transfection for Cas9 expression and subsequently single cell seeded at 0.2–1 cells per well in a 384 well plate. Single clones were pre-selected by lytic Nanoluc reconstitution under treatment with indisulam or DMSO. Clones that showed loss of luciferase signal under indisulam were subsequently characterized via immunoblotting for the introduced HiBit-tag and via genotyping by PCR of the targeted genomic region.

For generation of the RBM39 G268V mutant HCT116 cells, again annealed oligos were utilized in a similar fashion. After CRISPR/Cas9 mediated cutting at the specific genomic locus a repair template harboring a 40 bp homology and the desired point mutation in its center was used to introduce the mutation. Cells were seeded and transfected with the sgRNA/Cas9 plasmid and the repair oligos as mentioned above followed by pulse-selection for Cas9 expression from 24 h post transfection to 72 hrs post transfection and subsequently single cell seeding. Next, clones were mirror-plated in 96 well plates after initial expansion and pre-selected by treatment with indisulam in one of the mirror plates. Single clones that showed resistance to indisulam were subsequently characterized via genotyping by PCR of the targeted genomic region to identify the specific introduced point mutation.

### Transcript quantification via qPCR

1 x 10^6^ HEK293T cells with ectopic expression of HiBit-DCAF15 and LgBit were treated with DMSO or 10 μM indisulam for 12 h, detached and RNA was isolated using the RNeasy Kit and QIAshredder (Qiagen) following standard protocol with DNA digestion. Reverse transcription PCR was performed with the RevertAID First Strand cDNA synthesis kit and Oligo-dT primers (Thermo Scientific). DCAF15 RNA was quantified in a PCR reaction using SYBR select master mix (Fisher Scientific) in the following reaction: 3.75 μl of 1:10 diluted cDNA, 0.75 μl of DCAF15_exon9-11 primer mix (10 μM each, see Supplementary Table 1), 3 μl H2O and 7.5 μl SYBR master mix. The reaction mixture was denatured for 3’ at 95 °C followed by 45 cycles of 15” at 95 °C, 45” at 60 °C and 15” at 95 °C with a final extension of 1’ at 60 °C and 15” at 95 °C in a StepOne Plus real-time PCR cycler (Applied Biosystems). The cycle number for exponential amplification was determined and normalized to DMSO treated samples and visualized with Prism (GraphPad).

### Western blot analysis

PBS-washed cell pellets were lysed in RIPA Buffer (50 mM Tris-HCl pH 8.0, 150 mM NaCl, 1% Triton X-100, 0.5% sodium deoxycholate, 0.1% SDS, 1× Halt protease inhibitor cocktail, 25 U ml^−1^ Benzonase). Lysates were cleared by centrifugation for 15 min at 4 °C and 20,000*g*. Protein concentration was measured by BCA according to the manufacturer’s protocol (Thermo Scientific™ Pierce™ BCA Protein Assay Kit) and 4X LDS sample buffer was added. Proteins (20 μg) were separated on 4-12% SDS-PAGE gels and transferred to nitrocellulose membranes. Membranes were blocked with 5% milk in TBST for 30 min at RT. Primary antibodies were incubated in milk or TBST alone for 1 h at RT or 4 °C overnight. Secondary antibodies were incubated for 1 h at RT. Blots were developed with chemiluminescence films. Primary antibodies used: BRD4 (1:1000, Abcam, ab128874), BRD3 (1:1000, Bethyl Laboratories, A302-368A), BRD2 (1:1000, Bethyl Laboratories, A302-582A), SMARCA4 (1:1000, Bethyl Laboratories, A300-813A), SMARCA2 (1:1000, Cell Signaling Technology, #6889), cMYC (1:1000, Santa Cruz Biotechnology, sc-764), GSPT1 (1:1000, Abcam, ab49878), CDK9 (1:1000, Cell Signaling Technology, 2316S), CRBN (1:2000, kind gift of R. Eichner and F. Bassermann), VHL (1:1000, Cell Signaling Technology, 2738), ACTIN (1:5000, Sigma-Aldrich, A5441-.2ML), GAPDH (1:1000, Santa Cruz Biotechnology, sc-365062). Secondary antibodies used: Peroxidase-conjugated AffiniPure Goat Anti-Rabbit IgG (1:10000, Jackson ImmunoResearch, 111-035-003) and Peroxidase-conjugated AffiniPure Goat Anti-Mouse IgG (1:10000, Jackson ImmunoResearch, 115-035-003).

### E3 ligase luciferase measurements

#### Live cell measurements

For NanoLuc measurements, cells were diluted to 1 M cells ml^-1^ in media and 10 μl of this suspension seeded in a black 384-well plate (Corning, 3764). For large scale chemical screens, compounds were dispensed with an Echo 550 system and resuspended in 10 μl of media prior to cell seeding to obtain a final assay volume of 40 μl and compound concentrations of 10 μM. For small scale luciferase measurements, 10 μl of compound solution was added to each well with cell suspension. Positive (depending on assayed ligase) and negative (DMSO) control compounds were scattered over each plate to judge and eliminate plate positional effects. Finally, 20 μl of media supplemented with 50 mM HEPES (Sigma), 1:100 Endurazine Luciferase live cell substrate (Promega) and depending on condition, CSN5i-3 (MedChemExpress) were added to each well. Luciferase measurements were performed every 1 to 2 hours on an EnVision plate reader (PerkinElmer). Results were analysed by employing python (v3.8.5), pandas (v1.1.3) and numpy (v1.19.2) to normalize each well and timepoint per plate to its relative negative control measurement (DMSO) and depicted using matplotlib (v3.3.2) and seaborn (v0.11.0).

#### Lytic measurements

For Lytic endpoint measurements cells were seeded as mentioned above and NanoLuc abundance was determined via the Nano-Glo HiBit lytic detection kit (Promega) following manufacturers recommendations. Depending on the cell line used (HiBit- or NanoLuc tagged protein), LgBit was added to the final measurement mix or not. Results were analysed as described above and visualized with Prism (GraphPad).

### Time-resolved Förster resonance energy transfer

Protein constructs, expression and purification were performed as previously described^45^. Titrations of compounds in BODIPY FL-E7820 displacement assay were carried out by mixing 200 nM biotinylated Strep-II-Avi-tagged DCAF15 variants, 2 nM terbium-coupled streptavidin in assay buffer containing 50 mM Tris, pH 7.5, 200 mM NaCl, 0.1% Pluronic F-68 solution (Sigma) and 5 μM of BODIPY FL-E7820. After dispensing the assay mixture, an increasing concentration of small molecules was dispensed in the 384-well plate (Corning, 4514) using a D300e Digital Dispenser (HP) normalized to 2% DMSO and then incubated for 60 min at room temperature. After excitation of terbium fluorescence at 337 nm, emission at 490 nm (terbium) and 520 nm (BODIPY FL) were recorded with a 70-μs delay over 600 μs to reduce background fluorescence, and the reaction was followed over 10 cycles of each data point using a PHERAstar FSX microplate reader (BMG Labtech). The TR-FRET signal of each data point was extracted by calculating the 520/490 nm ratio. The IC50 values were estimated using the variable slope equation in Prism (GraphPad). All TR-FRET results are plotted as mean ± s.d. from three independent replicates (*n* = 3).

### Molecular Docking Analysis

The crystal structures of DCAF15-DDB1ΔB-DDA1 complex (PDB: 6Q0R) were prepared using the Protein Preparation Wizard in Maestro (Maestro release 2022-1, Epik version 5.9137). Default settings were used, except that all crystallographic water molecules > 5 Å from heteroatom groups were removed. The docking receptor grid was created using the Receptor Grid Generation module in Glide (Glide version 94137). The grid box and center were set to default by using the active site ligand (E7820), with the active site ligand excluded from the grid. The ligands were prepared using the LigPrep module with OPLS3 force field and default settings (LigPrep version 61137). The docking poses were generated using the LigandDocking protocol as implemented in Schrödinger Suite 2022-1. Default settings were used with the Standard Precision (SP) score function with flexible ligand sampling. Briefly, the grid box and center were set at default using the active site ligand, and no constraints were defined. The top pose with the lowest Glide SP score is shown for dCeMM5. Figures were generated in PyMOL (2.5.1, Schrödinger, LLC).

### Expression proteomics

Global proteomics were essentially performed as previously described.^34,35^ First, we compared overall proteome-wide changes in HAP1 WT cells treated with DMSO or CSN5i-3 (1mM and 250 nM, 8h). Second, we profiled dRRM-1 treatment (10 μM for 10 hrs) in HAP1 DCAF15^-/-^ cells overexpressing HiBit-DCAF15 and LgBit.

#### Sample preparation

30×10^6^ HAP1 cells per condition were collected, washed four times with ice-cold DPBS, the supernatant aspirated and pellets snapfrozen in liquid N2. Each washed cell pellet was lysed separately in 40 μL of freshly prepared lysis buffer containing 50 mM HEPES (pH 8.0), 2% SDS, 0.1 M DTT, 1 mM PMSF, and protease inhibitor cocktail (Sigma-Aldrich). Samples rested at RT for 20 minutes before heating to 99 °C for 5 min. After cooling down to RT, DNA was sheared by sonication using a Covaris S2 high performance ultrasonicator. Cell debris was removed by centrifugation at 20.000 g for 15 min at 20 °C. Supernatent was transferred to fresh eppendorf tubes and protein concentration determined using the BCA protein assay kit (Pierce Biotechnology). FASP was performed using a 30 kDa molecular weight cutoff filter (VIVACON 500; Sartorius Stedim Biotech) according to published procedures.^52^ In brief, 100 ug total protein per sample were reduced by adding DTT at a final concentration of 83.3 mM followed by incubation at 99 °C for 5 min. After cooling to room temperature, samples were mixed with 200 μL of freshly prepared 8 M urea in 100 mM Tris-HCl (pH 8.5) (UA-solution) in the filter unit and centrifuged at 14.000 g for 15 min at 20 °C to remove SDS. Any residual SDS was washed out with 200 μL of UA. The proteins were alkylated with 100 μL of 50 mM iodoacetamide in the dark for 30 min at RT. Next, three washing steps with 100 μL of UA solution were performed, followed by three washing steps with 100 μL of 50 mM TEAB buffer (Sigma-Aldrich). Proteins were digested with trypsin at a ratio of 1:50 overnight at 37 °C. Peptides were recovered using 40 μL of 50 mM TEAB buffer followed by 50 μL of 0.5 M NaCl (Sigma-Aldrich). Peptides were desalted using C18 solid phase extraction spin columns (The Nest Group). After desalting, peptides were labeled with TMT 10-plex reagents according to the manufacturer protocol (Pierce Biotechnology). After quenching of the labeling reaction, labeled peptides were pooled, organic solvent removed in vacuum concentrator and labeled peptides cleaned via C18 solid phase extraction.

#### Offline Fractionation via RP-HPLC at high pH

Tryptic peptides were re-buffered in 20 mM ammonium formiate buffer pH 10, shortly before separation by reversed phase liquid chromatography at pH 10 as described^53^. Peptides were separated into 96 time-based fractions on a column (150 x 2.0 mm Gemini-NX 3 μM C18 110 Å, Phenomenex) using an Agilent 1200 series HPLC system fitted with a binary pump delivering solvent at 100 uL/min. Acidified fractions were consolidated into 36 fractions via a concatenated strategy described^54^. After solvent removal in a vacuum concentrator, samples were reconstituted in 5% formic acid for LC-MS/MS analysis and kept at −80 °C until analysis.

#### 2D-RP/RP Liquid Chromatography Mass Spectrometry

Mass spectrometry was performed on an Orbitrap Fusion Lumos mass spectrometer (ThermoFisher Scientific) coupled to a Dionex Ultimate 3000RSLC nano system (ThermoFisher Scientific) via nanoflex source interface. Tryptic peptides were loaded onto a trap column (Pepmap 100, 5 μM, 5 x 0.3 mm, ThermoFisher Scientific) at a flow rate of 10 uL/min using 2% ACN and 0.05% TFA as loading buffer. After loading, the trap column was switched in-line with a 40 cm, 75 μM inner diameter analytical column (packed in-house with ReproSil-Pur 120 C18-AQ, 3 μM, Dr. Maisch). Mobile-phase A consisted of 0.4% formic acid in water and mobile-phase B of 0.4% formic acid in a mix of 90% acetonitrile and 9.6% water. The flow rate was set to 230 nL/min and a three-step 90 min gradient applied (6 to 30% solvent B within 81 min, 30 to 65% solvent B within 8 min, and 65 to 100% solvent B within 1 min, 100% solvent B for 6 min before equilibrating at 6% solvent B for 18 min prior to next injection). Analysis on the MS was performed in a data-dependent acquisition (DDA) mode using a maximum 3 s cycle time. Full MS^1^ scans were acquired in the Orbitrap with a scan range of 375 - 1650 m/z and a resolution of 120,000 (at 200 m/z). Automatic gain control (AGC) was set to a target of 2 x 10^5^ and a maximum injection time of 50 ms. MS^2^ spectra were acquired in the Orbitrap at a resolution of 50,000 (at 200 m/z) with a fixed first mass of 100 m/z. A tandem MS approach was chosen (TMT reporter ion intensities extracted from MS^2^ scans) to achieve maximum proteome coverage. To minimize TMT ratio compression effects by interference of contaminating co-eluting isobaric peptide ion species, precursor isolation width in the quadrupole was set to 0.4 Da and an extended fractionation scheme applied (36 fractions, see above). Monoisotopic peak determination was set to peptides with inclusion of charge states between 2 and 7. Intensity threshold for MS^2^ selection was set to 5 x 10^4^. Higher energy collision induced dissociation (HCD) was applied with a normalized collision energy (NCE) of 38%. AGC was set to 1 x 10^5^ with a maximum injection time of 105 ms. Dynamic exclusion for selected ions was 60 s. A single lock mass at m/z 445.120024 was employed. Xcalibur (v4.2.28.14) and Tune (v3.1, 2412.17) were used to operate the instrument.

#### Data Analysis

Acquired raw data files were processed using Proteome Discoverer (v2.2.0) with the Sequest HT database search engine and Percolator validation software node (v3.04) to remove false positives with a false discovery rate (FDR) of 1% on peptide and protein level under strict conditions. Searches were performed with full tryptic digestion against the human SwissProt database v2017.06 with up to two allowed miscleavage sites. Oxidation (+15.9949Da) of methionine was set as variable modification, while carbamidomethylation (+57.0214Da) of cysteine residues and TMT labeling of peptide N-termini and lysine residues were set as fixed modifications. Data was searched with mass tolerances of ± 10 ppm and ± 0.02Da on the precursor and fragment ions, respectively. Results were filtered to include peptide spectrum matches (PSMs) with Sequest HT cross-correlation factor (Xcorr) scores of ≥ 1 and high peptide confidence assigned by Percolator. MS^2^ signal-to-noise values (S/N) values of TMT reporter ions were used to estimate peptide/protein abundance changes. PSMs with precursor isolation interference values of ≥ 50% and average TMT-reporter ion S/N ≤ 10 were excluded from quantitation. Only unique peptides were used for TMT quantitation. Isotopic impurity correction and TMT channel normalization based on total peptide amount were applied. For statistical analysis *and P*-value calculation, the integrated ANOVA test was used. TMT ratios with *P*-values below 0.01 were considered as significant. Only proteins with > 1 peptide detected and > 1 unique peptide detected were considered for further analysis. A volcano plot was generated for all proteins detected by depicting log_2_ fold changes in TMT-ratios for CSN5i treatment vs. DMSO against the protein specific *P*-values. For visualizing dRRM-1 destabilized proteins a similar plot was generated with changes between degrader treatment and DMSO. Full results of global proteome changes can be found in Supplementary Table 2.

## Data availability

All data presented in this study is available in main figures, supplementary information or upon request from the authors. Full proteomics results are available in the supplementary information.

## Supplementary Figures

**Supplementary Figure 1.**
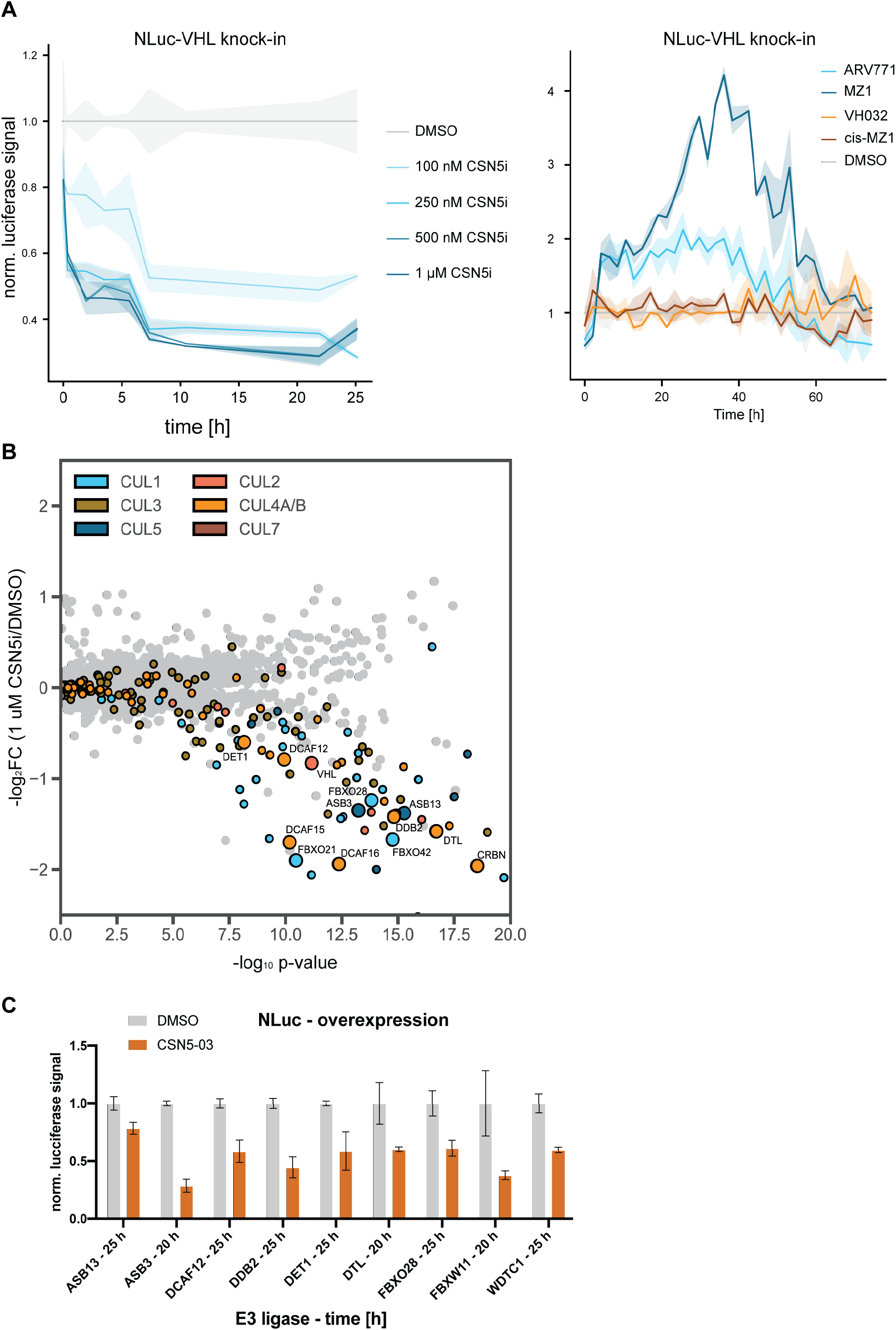
(A) DMSO normalized live-cell luciferase signal of HAP1 VHL-NanoLuc knock-in cells treated with different doses of CSN5i-3 or DMSO (left). On the right, same cells co-treated with CSN5i-3 (250 nM) and ARV-771, MZ-1, VH-032, cis-MZ1 or DMSO (1 μM each). Mean of *n* = 3 replicates. (B) Volcano plot depicting global log_2_-fold changes of protein abundance in HAP1 cells treated with CSN5i-3 (1 μM) for 8 h. CRL substrate receptors are labeled in the indicated colors. SRs selected for validation via luciferase tagging are highlighted. Data of *n* = 3 replicates. (C) DMSO normalized live-cell luciferase signal of HAP1 cells overexpressing the indicated protein in C-terminal fusion with NanoLuc. Cells were treated with DMSO, CSN5i-03 (500 nM) or CSN5i-03/MLN4924 (500 nM each) and measured at the indicated timepoint after treatment. Mean of n = 2 independent measurements. Representative data of *n* = 2 experiments.

**Supplementary Figure 2.**
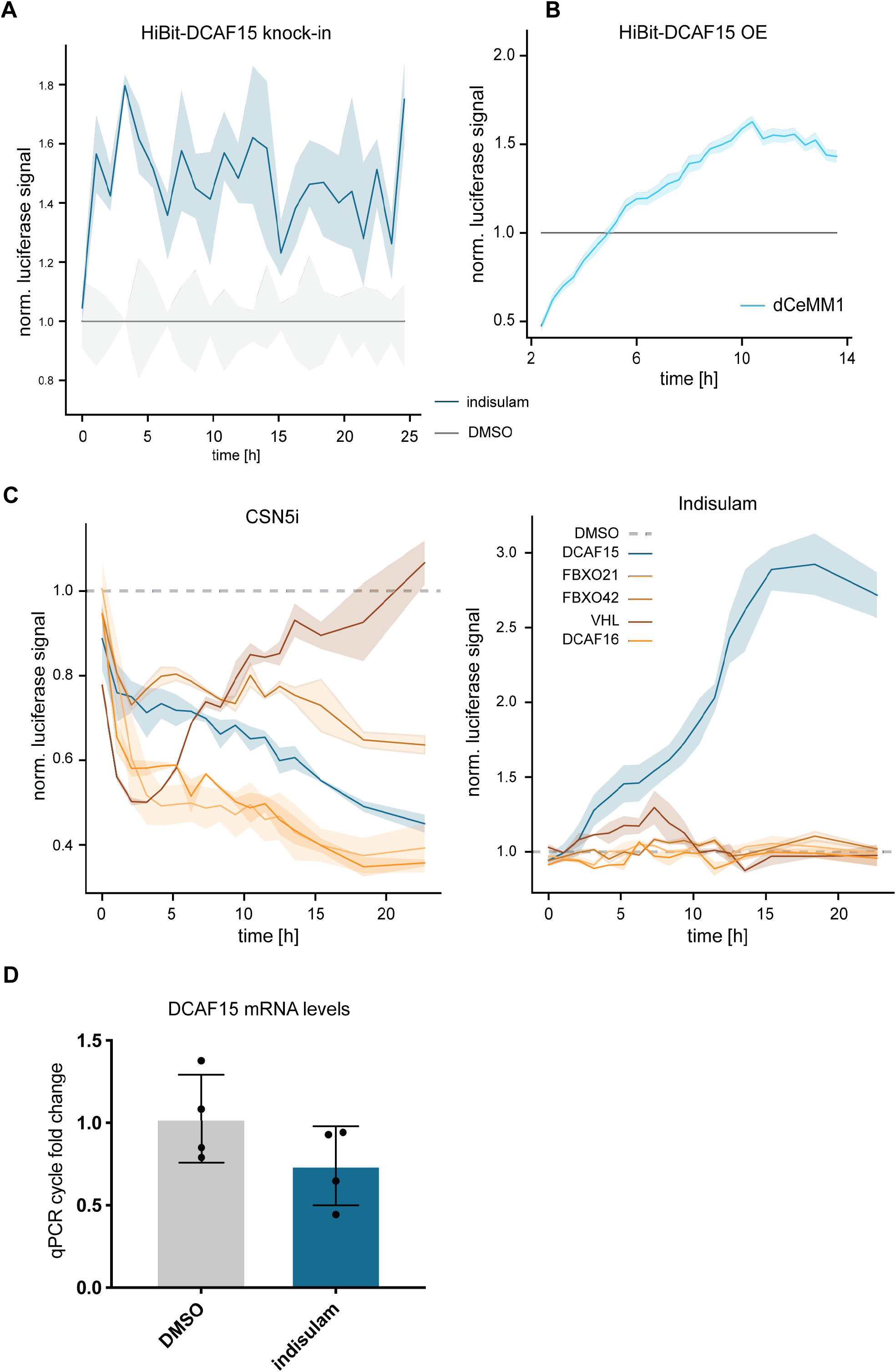
(A) DMSO normalized live-cell luciferase signal of HEK293t HiBit-DCAF15 endogenous knock-in cells with ectopic expression of LgBit treated with indisulam or DMSO (10 μM). Mean of *n* = 3 replicates. (B) DMSO normalized live-cell luciferase signal of HEK293t DCAF15^-/-^ cells with reconstitution of HiBit-DCAF15 + LgBit treated with dCeMM1 (10 μM) or DMSO. Mean of *n* = 3 replicates. Representative data of *n* = 3 experiments. (C) DMSO normalized live-cell luciferase signal of HAP1 cells with endogenous knock-in of NanoLuc for the indicated proteins. Cells were treated with DMSO, CSN5i-3 (250 nM, left) or CSN5i-3/indisulam (250 nM and 1 μM, right) and measured at the indicated timepoint after treatment. Mean of *n* = 2 replicates. Representative data of *n* = 2 experiments. (D) Bar graph depicting fold-change in qPCR cycles of exponential amplification in HEK293t DCAF15^-/-^ cells with reconstitution of HiBit-DCAF15 + LgBit after treatment with indisulam or DMSO for 10 h. Mean of *n* = 4 replicates.

**Supplementary Figure 3.**
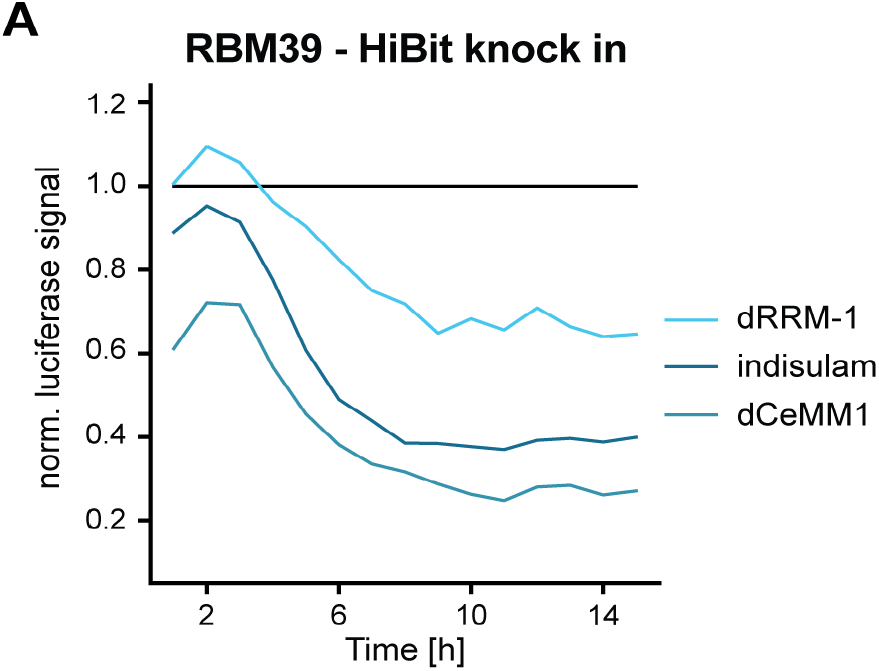
(A) DMSO normalized live-cell luciferase signal of HCT116 RBM39-HiBit knock in cells with ectopic expression of LgBit treated with indisulam, dCeMM1, dRRM-1 or DMSO (10 μM each).

## References

1. Stanton, B. Z., Chory, E. J. & Crabtree, G. R. Chemically induced proximity in biology and medicine. Science 359, eaao5902 (2018).

2. Gerry, C. J. & Schreiber, S. L. Unifying principles of bifunctional, proximity-inducing small molecules. Nat Chem Biol 16, 369–378 (2020).

3. Deshaies, R. J. Multispecific drugs herald a new era of biopharmaceutical innovation. Nature 580, 329–338 (2020).

4. Poirson, J. et al. Proteome-scale induced proximity screens reveal highly potent protein degraders and stabilizers. bioRxiv (2022).

5. Chamberlain, P. P. & Hamann, L. G. Development of targeted protein degradation therapeutics. Nature chemical biology 15, 937–944 (2019).

6. Burslem, G. M. & Crews, C. M. Proteolysis-Targeting Chimeras as Therapeutics and Tools for Biological Discovery. Cell 181, 102–114 (2020).

7. Jevtić, P., Haakonsen, D. L. & Rapé, M. An E3 ligase guide to the galaxy of small-molecule-induced protein degradation. Cell Chem Biol 28, 1000–1013 (2021).

8. Petroski, M. D. & Deshaies, R. J. Function and regulation of cullin-RING ubiquitin ligases. Nat Rev Mol Cell Biol 6, 9–20 (2005).

9. Harper, J. W. & Schulman, B. A. Cullin-RING Ubiquitin Ligase Regulatory Circuits: A Quarter Century Beyond the F-Box Hypothesis. Annu Rev Biochem 90, 403–429 (2021).

10. Reitsma, J. M. et al. Composition and Regulation of the Cellular Repertoire of SCF Ubiquitin Ligases. Cell 171, 1326–1339.e14 (2017).

11. Reichermeier, K. M. et al. PIKES Analysis Reveals Response to Degraders and Key Regulatory Mechanisms of the CRL4 Network. Mol Cell 77, 1092–1106.e9 (2020).

12. Pierce, N. W. et al. Cand1 promotes assembly of new SCF complexes through dynamic exchange of F box proteins. Cell 153, 206–215 (2013).

13. Baek, K. et al. NEDD8 nucleates a multivalent cullin-RING-UBE2D ubiquitin ligation assembly. Nature 578, 461–466 (2020).

14. Cope, G. A. et al. Role of predicted metalloprotease motif of Jab1/Csn5 in cleavage of Nedd8 from Cul1. Science 298, 608–611 (2002).

15. Cavadini, S. et al. Cullin-RING ubiquitin E3 ligase regulation by the COP9 signalosome. Nature 531, 598–603 (2016).

16. Galan, J. M. & Peter, M. Ubiquitin-dependent degradation of multiple F-box proteins by an autocatalytic mechanism. Proc Natl Acad Sci U S A 96, 9124–9129 (1999).

17. Zhou, P. & Howley, P. M. Ubiquitination and degradation of the substrate recognition subunits of SCF ubiquitin-protein ligases. Mol Cell 2, 571–580 (1998).

18. Békés, M., Langley, D. R. & Crews, C. M. PROTAC targeted protein degraders: the past is prologue. Nat Rev Drug Discov 21, 181–200 (2022).

19. Lu, G. et al. The myeloma drug lenalidomide promotes the cereblon-dependent destruction of Ikaros proteins. Science 343, 305–309 (2014).

20. Krönke, J. et al. Lenalidomide causes selective degradation of IKZF1 and IKZF3 in multiple myeloma cells. Science 343, 301–305 (2014).

21. Sievers, Q. L. et al. Defining the human C2H2 zinc finger degrome targeted by thalidomide analogs through CRBN. Science 362, eaat0572 (2018).

22. Han, T. et al. Anticancer sulfonamides target splicing by inducing RBM39 degradation via recruitment to DCAF15. Science 356, eaal3755 (2017).

23. Uehara, T. et al. Selective degradation of splicing factor CAPERα by anticancer sulfonamides. Nat Chem Biol 13, 675–680 (2017).

24. Matyskiela, M. E. et al. A novel cereblon modulator recruits GSPT1 to the CRL4(CRBN) ubiquitin ligase. Nature 535, 252–257 (2016).

25. Henning, N. J. et al. Discovery of a Covalent FEM1B Recruiter for Targeted Protein Degradation Applications. J Am Chem Soc 144, 701–708 (2022).

26. Spradlin, J. N. et al. Harnessing the anti-cancer natural product nimbolide for targeted protein degradation. Nat Chem Biol 15, 747–755 (2019).

27. Luo, M. et al. Chemoproteomics-enabled discovery of covalent RNF114-based degraders that mimic natural product function. Cell Chem Biol 28, 559–566.e15 (2021).

28. Zhang, X., Crowley, V. M., Wucherpfennig, T. G., Dix, M. M. & Cravatt, B. F. Electrophilic PROTACs that degrade nuclear proteins by engaging DCAF16. Nat Chem Biol 15, 737–746 (2019).

29. Zhang, X. et al. DCAF11 Supports Targeted Protein Degradation by Electrophilic Proteolysis-Targeting Chimeras. J Am Chem Soc 143, 5141–5149 (2021).

30. Simonetta, K. R. et al. Prospective discovery of small molecule enhancers of an E3 ligase-substrate interaction. Nat Commun 10, 1402 (2019).

31. Słabicki, M. et al. The CDK inhibitor CR8 acts as a molecular glue degrader that depletes cyclin K. Nature 585, 293–297 (2020).

32. Mayor-Ruiz, C. et al. Rational discovery of molecular glue degraders via scalable chemical profiling. Nat Chem Biol 16, 1199–1207 (2020).

33. Cope, G. A. & Deshaies, R. J. COP9 signalosome: a multifunctional regulator of SCF and other cullin-based ubiquitin ligases. Cell 114, 663–671 (2003).

34. Schlierf, A. et al. Targeted inhibition of the COP9 signalosome for treatment of cancer. Nat Commun 7, 13166 (2016).

35. Mayor-Ruiz, C. et al. Plasticity of the Cullin-RING Ligase Repertoire Shapes Sensitivity to Ligand-Induced Protein Degradation. Molecular Cell 75, 849–858.e8 (2019).

36. Zengerle, M., Chan, K. H. & Ciulli, A. Selective Small Molecule Induced Degradation of the BET Bromodomain Protein BRD4. ACS Chem Biol 10, 1770–1777 (2015).

37. Bondeson, D. P. et al. Catalytic in vivo protein knockdown by small-molecule PROTACs. Nat Chem Biol 11, 611–617 (2015).

38. Raina, K. et al. PROTAC-induced BET protein degradation as a therapy for castration-resistant prostate cancer. Proc Natl Acad Sci U S A 113, 7124–7129 (2016).

39. Gadd, M. S. et al. Structural basis of PROTAC cooperative recognition for selective protein degradation. Nat Chem Biol 13, 514–521 (2017).

40. Farnaby, W. et al. BAF complex vulnerabilities in cancer demonstrated via structure-based PROTAC design. Nat Chem Biol 15, 672–680 (2019).

41. Riching, K. M. et al. Quantitative Live-Cell Kinetic Degradation and Mechanistic Profiling of PROTAC Mode of Action. ACS Chem Biol 13, 2758–2770 (2018).

42. Riching, K. M., Mahan, S. D., Urh, M. & Daniels, D. L. High-Throughput Cellular Profiling of Targeted Protein Degradation Compounds using HiBiT CRISPR Cell Lines. J Vis Exp (2020).

43. Bussiere, D. E. et al. Structural basis of indisulam-mediated RBM39 recruitment to DCAF15 E3 ligase complex. Nat Chem Biol 16, 15–23 (2020).

44. Du, X. et al. Structural Basis and Kinetic Pathway of RBM39 Recruitment to DCAF15 by a Sulfonamide Molecular Glue E7820. Structure 27, 1625–1633.e3 (2019).

45. Faust, T. B. et al. Structural complementarity facilitates E7820-mediated degradation of RBM39 by DCAF15. Nature Chemical Biology 16, 7–14 (2020).

46. Koduri, V. et al. Targeting oncoproteins with a positive selection assay for protein degraders. Sci Adv 7, eabd6263 (2021).

47. Ting, T. C. et al. Aryl Sulfonamides Degrade RBM39 and RBM23 by Recruitment to CRL4-DCAF15. Cell Rep 29, 1499–1510.e6 (2019).

48. Hanzl, A. et al. Charting functional E3 ligase hotspots and resistance mechanisms to small-molecule degraders. bioRxiv (2022).

49. Yang, X. et al. A public genome-scale lentiviral expression library of human ORFs. Nat Methods 8, 659–661 (2011).

50. Jaeger, M. G. et al. Selective Mediator dependence of cell-type-specifying transcription. Nat Genet 52, 719–727 (2020).

51. Brand, M. & Winter, G. E. Locus-Specific Knock-In of a Degradable Tag for Target Validation Studies. Methods Mol Biol 1953, 105–119 (2019).

52. Wisniewski, J. R., Zougman, A., Nagaraj, N. & Mann, M. Universal sample preparation method for proteome analysis. Nat Methods 6, 359–362 (2009).

53. Gilar, M., Olivova, P., Daly, A. E. & Gebler, J. C. Two-dimensional separation of peptides using RP-RP-HPLC system with different pH in first and second separation dimensions. J Sep Sci 28, 1694–1703 (2005).

54. Wang, Y. et al. Reversed-phase chromatography with multiple fraction concatenation strategy for proteome profiling of human MCF10A cells. Proteomics 11, 2019–2026 (2011).

